# A cross-tissue POSTN+ fibroblast atlas links periodontal, tumor, and fibrotic stromal niches

**DOI:** 10.64898/2026.05.20.726414

**Authors:** Chengze Wang, Haiping Yang, Mengna Lin, Ying Wang, Guoli Yang

**Affiliations:** Stomatology Hospital, School of Stomatology, Zhejiang University School of Medicine, Clinical Research Center for Oral Diseases of Zhejiang Province, Key Laboratory of Oral Biomedical Research of Zhejiang Province, Cancer Center of Zhejiang University, Hangzhou 310006, China

**Keywords:** fibroblast, POSTN, periostin, cross-tissue atlas, single-cell RNA-seq, cancer-associated fibroblasts, periodontium, liver fibrosis, KLF4

## Abstract

Cross-tissue single-cell atlases have re-framed fibroblasts as a continuum of activation states, with universal Pi16+ progenitors giving rise to tissue-restricted activated populations[1] and shared pathological states recurring across inflammatory diseases[2]. Periostin (POSTN), a matricellular protein of injured, fibrotic, and tumor stroma, has been independently linked to activated fibroblasts in liver fibrosis[3], colorectal cancer[4], head-and-neck cancer[5], and dental contexts[6, 7], but cross-tissue conservation of a single POSTN+ program is untested. Here we built a Harmony-integrated atlas of 56,713 human and mouse fibroblasts from eight single-cell datasets spanning six organ contexts (periodontal ligament, periodontitis, oral squamous-cell carcinoma, colorectal cancer, temporomandibular-joint osteoarthritis, and bile-duct-ligation liver fibrosis). A conservative cluster-consensus definition (Wilcoxon padj < 0.05 and log2FC > 0.5 within an atlas-integrated leiden cluster, combined with per-cell POSTN > 0) identified 11,451 POSTN+ cells (20.2% of the atlas) recurring across all six contexts at frequencies from 6.2% (periodontal ligament) to 55.1% (liver fibrosis). Within-fibroblast differential expression yielded a 102-gene shared core program — collagen biosynthesis, ECM crosslinkers, and matricellular markers including POSTN, SPARC, BGN, FN1, MMP2, and CTHRC1 — interpreted as POSTN-specific transcriptional amplification of an activated ECM-remodelling module. KLF4, hypothesized a priori as a POSTN+ co-marker, was upregulated in only one of six contexts, consistent with its role as a quiescence brake released during activation[3]. Three pre-registered sensitivity analyses (Harmony parameter, three definitions, dataset exclusion) and an independent Puram-2017 OSCC cohort (1,422 fibroblasts; 101/102 core genes recovered; primary vs lymph-node-met Mann–Whitney p = 0.005) support robustness across integration parameters, definitions, dataset inclusion, and platform.

## Introduction

Fibroblasts have historically functioned as a residual diagnosis — the cells left over once epithelium, endothelium, and immune populations have been gated out — and the past decade of single-cell transcriptomics has shown how misleading that framing is[8, 9]. Across organs, fibroblasts span a continuum of states whose boundaries are set by tissue context, mechanical environment, and inflammatory cues, not by a single canonical marker. Two cross-tissue atlases have anchored this re-framing. Buechler and colleagues integrated mouse fibroblasts from steady-state tissues and identified a universal Pi16+ progenitor subset that gives rise to tissue-restricted activated populations[1]. Korsunsky and colleagues subsequently atlased pathological fibroblasts in rheumatoid arthritis, inflammatory bowel disease, idiopathic pulmonary fibrosis, and systemic sclerosis, showing that disease-associated activation states recur across organs[2]. Both works converge on the idea that activation programs, rather than lineage labels, define what a fibroblast does in disease.

Within this re-framed view, POSTN (periostin) has emerged as a candidate cross-tissue activation marker. POSTN encodes a matricellular protein induced in repair, fibrosis, and cancer-associated stroma[10], and POSTN+ fibroblasts have been described independently in liver fibrosis as the activated derivative of a Clec3b+ portal-fibroblast state held in check by KLF4[3], in human colorectal cancer as one of eleven CAF subtypes[4], in head-and-neck squamous-cell carcinoma as a matrix-remodelling stromal population[5], and in periodontal-ligament[6], temporomandibular-joint[7, 11], and gingival inflammation[12] contexts. Each of these reports frames POSTN locally, however, and no integrated test has asked whether the same POSTN+ activation program recurs across these tissues, what its conserved transcriptional core consists of, or how it relates to the KLF4-marked quiescent baseline. The Buechler and Korsunsky atlases do not include oral or dental fibroblasts, did not focus on the POSTN axis, and treat tumor-associated fibroblasts only marginally; the question of whether POSTN+ defines a single activation state across mechanically loaded periodontal niches, tumor stroma, and chronic fibrosis is therefore open.

We addressed this question by extending the cross-tissue fibroblast framework along the POSTN axis. Eight single-cell datasets covering six organ contexts — periodontal ligament[6], periodontitis (two cohorts[12, 13]), oral squamous-cell carcinoma[14], colorectal cancer[4], temporomandibular-joint osteoarthritis (two cohorts[7, 11]), and bile-duct-ligation liver fibrosis[3] — were harmonized to a common human-symbol gene space, integrated with Harmony[15], and probed for a POSTN+ activation module. Our pre-registered analysis plan defines POSTN+ as a conservative cluster-consensus state, derives a within-fibroblast shared core program, and exposes the POSTN+/KLF4 quiescence-vs-activation axis. We further pre-registered three sensitivity analyses — Harmony parameter θ, three competing POSTN+ definitions, and exclusion of the only dataset distributed without raw counts — together with an independent validation cohort (Puram 2017 OSCC[5]) used outside the integration. The atlas, the 102-gene shared core program, and the sensitivity tables are provided as a per-figure JSON sidecar registry against which every cited number in this manuscript is auditable.

## Results

### A Harmony-integrated atlas of 56,713 fibroblasts spans six organ contexts

We assembled eight single-cell datasets covering six organ contexts (Fig. 1A; Methods). Per-dataset preprocessing respected each cohort’s published cell-type assignments where available; mouse data (temporomandibular-joint cohorts and the liver bile-duct-ligation cohort) were re-indexed to human gene symbols using a 1:1 ortholog map derived from MGI HOM_MouseHumanSequence (18,782 unique 1:1 pairs retained from 21,930 mouse and 24,592 human entries). The eight datasets were concatenated on the intersection of expressed genes (14,545 genes; 14,296 retained after a min-cells = 10 filter), normalized to a target sum of 10^4^ counts per cell with log1p, and reduced by PCA on the full filtered gene set within scanpy[16]. Harmony[15] was applied with batch_key = dataset_alias and converged in five iterations under default parameters (θ = 2, σ = 0.1, max_iter = 20). The Harmony embedding produced a UMAP in which biological replicates within tissue contexts mixed coherently while contexts retained globally distinct positions (Fig. 1A), and a leiden[17] clustering at resolution 0.5 (igraph backend, two iterations, undirected graph) yielded 56 clusters that we used as the substrate for the POSTN+ definition. The mean cluster-level dataset entropy of the integrated atlas was 0.303, against a theoretical maximum of 3.000 for eight batches. Although low entropy can in principle reflect under-correction, two robustness analyses indicate that the structure we report is not an artefact of the default Harmony parameterization. First, a parameter-invariance analysis (Fig. 1C) shows the POSTN+ structure is preserved under a more aggressive Harmony parameterization (θ = 4) that nearly doubles the entropy (0.303 → 0.593). Second, a method-matched comparison against scVI[18] — restricted to the seven datasets distributed with raw counts (the eighth dataset, perio_mmp_gse171213_caf, is shipped only as scaled values and is incompatible with scVI’s negative-binomial likelihood; periodontitis context remains covered by perio_byrd2024) — recovers comparable batch-mixing entropy under both Harmony at matched input (0.860, 30.6% of the seven-batch theoretical maximum of log2(7) = 2.807) and scVI (0.949, 33.8%), substantially above the 0.303 of the primary eight-dataset run. The eight-dataset entropy is therefore primarily input-driven (the scaled perio_mmp_gse171213_caf cohort clusters apart from the rest) rather than reflecting Harmony-specific under-correction, and per-cell POSTN expression positivity is method-invariant across all six contexts (Fig. S3D). A full decomposition of how integration choices interact with the cluster-consensus POSTN+ definition is given in the Limitations.

**Figure 1.**
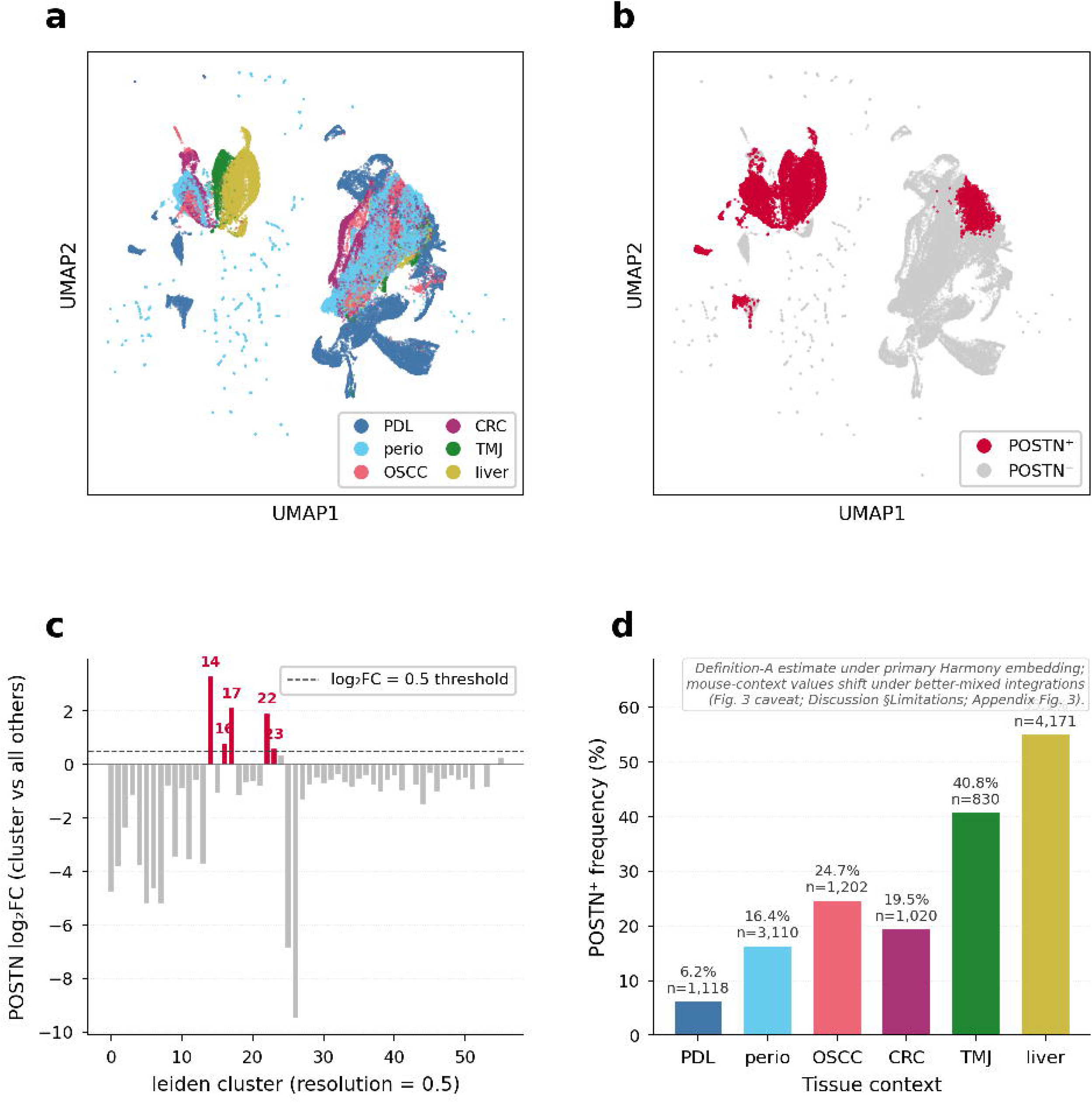
POSTN+ pan-fibroblast atlas across six organ contexts (4-block composite). A. Atlas construction. Harmony-integrated UMAP of 56,713 fibroblasts from eight single-cell datasets covering six organ contexts (PDL, perio, OSCC, CRC, TMJ, liver), coloured by tissue context (left) and dataset (right); 50 PCs, 56 leiden clusters at resolution 0.5. **B. POSTN+ cluster-consensus definition**. Left: UMAP highlighting 11,451 POSTN+ cells (20.2% of atlas; red on grey) defined as cells in candidate clusters meeting Wilcoxon padj < 0.05 and log2FC > 0.5 with non-zero per-cell POSTN expression. Middle: per-cluster POSTN log2FC against all other clusters; the five candidate clusters (14, 16, 17, 22, 23) are highlighted in red and exceed the dashed log2FC = 0.5 threshold. Right: per-context POSTN+ frequency under the primary 8-set Harmony embedding, ranging from 6.2% in periodontal ligament to 55.1% in liver fibrosis. These per-context percentages are definition-A estimates under the primary Harmony embedding, not integration-invariant biological prevalences — the mouse-context values in particular shift substantially under better-mixed embeddings because the cluster-consensus filter interacts with how mouse cells are clustered (Fig. S3 and Discussion Section Limitations); per-cell POSTN expression positivity is method-invariant. **C. Robustness**. Per-context POSTN+ frequency at default Harmony θ = 2 versus aggressive θ = 4; per-context rank Spearman ρ = 1.000 (n = 6); maximum absolute shift 4.4 percentage points. **D. Shared POSTN+ core program**. Heatmap of the top fifty (of 102) genes upregulated within fibroblasts in POSTN+ versus POSTN-cells in at least four of six contexts; rows = genes ranked by mean cross-context log2FC, columns = contexts, values = log2((mean POSTN+ + 1) / (mean POSTN-+ 1)) per context; * marks expected ECM and POSTN-program markers; POSTN itself is up in 6/6 contexts.

### POSTN+ fibroblasts recur across all six organ contexts under a conservative cluster-consensus definition

We pre-registered a conservative two-condition definition for POSTN+ status (Methods). At the cluster level, we required that the cell’s atlas-integrated leiden cluster meet Wilcoxon padj < 0.05 and log2FC > 0.5 (computed as log2((mean_in + 1)/(mean_out + 1)) on raw counts) for POSTN against all other clusters; five clusters (14, 16, 17, 22, and 23) met this criterion. At the cell level, we required that the cell’s normalized POSTN expression be greater than zero. Cells satisfying both conditions were classified POSTN+. This definition is conservative by design: high-POSTN expressors that fall outside the candidate clusters are filtered out, trading sensitivity for specificity and ensuring that POSTN+ cells inhabit POSTN-enriched neighborhoods.

Under this definition, 11,451 cells (20.2% of the atlas) were classified POSTN+ (Fig. 1B). All six organ contexts contributed. Per-context POSTN+ frequencies under definition A spanned an order of magnitude — periodontal ligament 6.2%, periodontitis 16.4%, colorectal cancer 19.5%, oral squamous-cell carcinoma 24.7%, temporomandibular-joint osteoarthritis 40.8%, and liver fibrosis 55.1% (Fig. 1B, right panel). These per-context numbers should be read as definition-A estimates rather than stable biological prevalences, because broader expression-based definitions B and C shift the absolute frequencies materially in some contexts (most visibly periodontitis and OSCC; Table S1). What is robust across definitions is the cross-context ordering — contexts dominated by active fibrotic remodelling or tumor stroma harbor the largest POSTN+ pools and resting periodontal ligament the smallest — and the recurrence of POSTN+ across all six contexts under all sensitivity perturbations described below.

### A 102-gene shared core program defines POSTN+ activation across contexts

To characterize what POSTN+ cells share transcriptionally across tissues, we performed within-fibroblast differential expression on raw counts in each context, comparing POSTN+ to POSTN-cells of the same context, and aggregated genes that crossed log2FC > 1 with mean POSTN+ counts ≥ 0.5 in at least four of six contexts. This produced a 102-gene shared core program (Fig. 1D). It is essential to emphasize the comparison structure: because the differential expression is POSTN+ versus POSTN-within fibroblasts, the program is not a generic fibroblast signature relative to other cell types but a within-fibroblast amplification programme that distinguishes activated POSTN+ cells from their POSTN-counterparts.

The 102-gene set is dominated by extracellular-matrix biosynthesis and remodelling. POSTN itself was upregulated in 6/6 contexts (per-context log2FC ranging from +2.90 in periodontitis to +6.11 in periodontal ligament), and the top fifty genes ranked by mean cross-context log2FC contained eleven collagen chains (COL1A1, COL1A2, COL3A1, COL5A1, COL5A2, COL6A1, COL6A2, COL6A3, COL11A1, COL12A1, and others), the matricellular markers SPARC and BGN, the fibronectin master gene FN1, the matrix-metalloproteinase / inhibitor pair MMP2 / TIMP1, the procollagen chaperone SERPINH1, the lysyl-oxidase-like crosslinking machinery, and the activated-fibroblast markers CTHRC1, AEBP1, ASPN, and CALD1 (Fig. 1D). We frame this gene set not as the discovery of an unexpected program but as evidence that POSTN+ cells transcriptionally amplify an activated ECM-remodelling module that POSTN-fibroblasts already express at baseline; the novelty resides in demonstrating that this amplification is conserved across periodontal, oral, joint, hepatic, and tumor stromal contexts within a single integrated atlas.

### KLF4 marks quiescence rather than POSTN+ activation, recovering the Clec3b portal-fibroblast model cross-tissue

Our project’s a-priori hypothesis listed KLF4 as a candidate POSTN+ co-marker. The data did not support this prediction. Of the six organ contexts, KLF4 was upregulated in POSTN+ versus POSTN-cells in only one (periodontal ligament); in the remaining five contexts, KLF4 was either non-significant or downregulated, and KLF4 did not enter the 102-gene shared core program. We initially treated this as a negative result, but the pattern recovers an explicit mechanistic prediction. In bile-duct-ligation liver fibrosis, Clec3b+ portal fibroblast activation is governed by a KLF4/periostin axis: KLF4 directly binds the *POSTN* promoter and represses its transcription, while POSTN, acting via αvβ5 integrin, drives portal fibroblast activation and ECM remodelling — so loss of KLF4 is the proximate event that permits POSTN induction[3]. Our cross-tissue data are compatible with this transcriptional-repression model beyond the liver: POSTN+ tracks the activated state and KLF4 tracks the quiescent baseline at the transcriptional level, suggesting a reciprocal quiescence-versus-activation pattern that may recur across tissues. This reframes our original hypothesis from a co-expression model to a candidate reciprocal-regulation model that remains to be tested functionally outside the liver, and the observation is consistent — rather than in conflict — with the prior literature.

### POSTN+ structure is robust to integration parameter, definition choice, and dataset inclusion

We pre-registered three robustness analyses. First, we re-integrated the same eight datasets with Harmony at θ = 4, σ = 0.05, max_iter = 30 — substantially more aggressive batch correction than the default (Fig. 1C). The mean cluster-level dataset entropy nearly doubled (0.303 → 0.593), indicating that the default parameterization is somewhat under-correcting; nevertheless, every per-context POSTN+ frequency shifted by less than 4.4 percentage points, and the per-context ranking was perfectly preserved (Spearman ρ = 1.000, n = 6 contexts; the rank Spearman is reported on n = 6 with the understanding that the exact permutation test under the null of independent rankings has p = 0.0028 and that we therefore avoid asymptotic p = 0 reporting). The set of candidate clusters expanded from five (default θ) to six (θ = 4), the total POSTN+ pool shifted from 11,451 to 10,608 cells, and the top-25 shared core genes overlapped 18/25 (72%) between parameterizations. We interpret this not as evidence that “low entropy is biological” but specifically as parameter-invariance of the POSTN+ structure within the Harmony family — a weaker and more defensible claim.

Second, we evaluated three competing POSTN+ definitions on the same atlas: the conservative cluster-consensus definition (A; n = 11,451, 20.2%), a percentile threshold (B; ≥ 70th-percentile raw POSTN counts with at least one read; n = 17,948, 31.6%), and a six-gene module score (C; POSTN + COL1A1 + SPARC + BGN + FN1 + TGFB1 in the top 30%; n = 17,014, 30.0%). The conservative definition was a strict subset of the broader expression-based pools: 85.4% of definition-A cells were also captured by B, and 71.2% by C. Per-context POSTN+ frequency rankings were Spearman-correlated at ρ = 0.71 (A vs B) and ρ = 0.89 (A vs C); B and C were correlated at ρ = 0.94 (n = 6, exact permutation). The largest discrepancy occurred in periodontitis, where definition A returns 16.4% but B and C return ≈ 35–41%; this reflects the design feature of the cluster-consensus filter, which excludes high-POSTN expressors that lie outside the candidate clusters. We retain definition A as primary because its outputs are auditable, rank-consistent with broader pools, and well-anchored to the atlas-integrated cluster structure; full pairwise overlap, per-context frequencies, and triple-agreement counts under all three definitions are tabulated in Table S1.

Third, we excluded the periodontitis MMP-CAF cohort (perio_mmp_gse171213_caf, n = 1,961), the only dataset distributed without raw counts; this re-derivation produced 105 shared core genes, of which 100 of the original 102 were retained (98% overlap). All twenty-five a-priori expected ECM markers were preserved, including POSTN itself (still 6/6 contexts), the eleven collagen chains, SPARC, BGN, FN1, MMP2, TIMP1, AEBP1, CTHRC1, CALD1, and SERPINH1. We conclude that the normalized-only cohort does not drive the shared core program.

Fourth, to ask whether the cross-species ortholog mapping (mouse → human via 1:1 MGI HOM table) is required for the shared-core signal, we removed all three mouse cohorts (tmj_shu_fib_subset, tmj_jci_gse267942, liver_clec3b; 9,607 cells) and re-derived the shared core from the four remaining human contexts (PDL, perio, OSCC, CRC; 47,106 cells, 5 datasets) at a parallel ≥ 3-of-4 (75%) threshold. **All 102 baseline shared-core genes were recoverable from the human contexts alone** (Table S3); POSTN itself was upregulated in 4/4 human contexts (per-context log2FC: PDL 4.75, perio 2.40, CRC 3.14, OSCC 2.80), as were all twenty-five top expected ECM markers (POSTN, eleven collagen chains, SPARC, BGN, FN1, MMP2, TIMP1, COL11A1, COL12A1, AEBP1, CTHRC1, CALD1, LGALS1, VCAN, ASPN, SERPINH1, TPM4 — TIMP1 and ASPN at 3/4, the rest at 4/4). We note that the human-only re-derivation identified 238 shared genes in total because the 3-of-4 (75%) cross-context cut admits more genes than the original 4-of-6 (67%) cut and does not constitute a stricter test; this analysis therefore demonstrates that the baseline program is recoverable in the human contexts alone, not that mouse cohorts contribute no species-specific signal. The mouse contexts (TMJ, liver) provide convergent rather than necessary evidence for the cross-tissue conservation claim.

### Independent validation in the Puram-2017 OSCC cohort

To verify that the atlas-derived 102-gene core program is not an artefact of the integration choices we made, we reproduced the analysis in an independent OSCC cohort that we did not include in the main atlas. Puram and colleagues’ GSE103322 cohort is distributed as TPM rather than raw counts and is therefore not directly compatible with the raw-count-aware Harmony pipeline; we used it as a supplementary validation cohort outside integration. Of 5,902 cells in the GSE103322 matrix, 1,422 were annotated as fibroblasts by the original authors, of which 1,112 were associated with primary tumours and 310 with lymph-node metastases. Of the 102 atlas-derived shared core genes, 101 (99%) were present in the Puram TPM matrix. Per-cell scores for the 102-gene program (computed by sc.tl.score_genes) reproduced the pattern observed in the integrated atlas, and were significantly higher in lymph-node-metastatic fibroblasts (mean = 2.85; n = 310) than in primary-tumour fibroblasts (mean = 2.76; n = 1,112; Mann–Whitney U p = 0.0046; Welch’s t-test p = 0.049; Fig. S1). The atlas-derived program therefore reproduces in an independent cohort acquired on a different platform, and tracks with metastatic site within OSCC.

## Discussion

We integrated eight single-cell fibroblast datasets covering six organ contexts and tested whether a single POSTN+ activation program recurs across them. The data support that interpretation. POSTN+ fibroblasts, defined conservatively as cluster-consensus high-POSTN expressors, were present in all six contexts at frequencies spanning an order of magnitude; the 102-gene shared core program was dominated by collagen biosynthesis, ECM crosslinking, and matricellular markers; KLF4, hypothesized a priori as a co-marker, instead behaved as a quiescence brake released during activation, recovering the Clec3b portal-fibroblast model[3] cross-tissue. Three pre-registered sensitivity analyses and an independent OSCC validation cohort support robustness of this structure to integration parameters, definition choice, dataset inclusion, and platform; a method-matched scVI comparison (Fig. S3) additionally decomposes the per-context POSTN+ frequencies into a method-invariant per-cell expression signal and a method-dependent cluster-consensus filter, so that the load-bearing claims do not depend on absolute mouse-context POSTN+ percentages.

Our work extends rather than displaces the existing pan-fibroblast frameworks. Buechler and colleagues established a steady-state mouse cross-tissue lineage including a universal Pi16+ progenitor pool[1]; Korsunsky and colleagues atlased pathological fibroblast states across four inflammatory diseases[2]. Neither framework includes oral, dental, or temporomandibular-joint fibroblasts; neither focuses on the POSTN axis; and tumor stroma is treated only marginally. To our knowledge no prior pan-fibroblast atlas has explicitly tested the conservation of a single POSTN-amplified activation program across more than three tissue contexts simultaneously. Our contribution is to do so for the POSTN axis specifically across human disease contexts, oral and dental tissues, tumor-associated fibroblasts (oral squamous-cell carcinoma and colorectal cancer), and chronic liver fibrosis, and to show that the within-fibroblast POSTN+ amplification of an activated ECM-remodelling module is conserved across these contexts. The novelty is the cross-tissue recurrence of the POSTN-amplified activated programme together with the KLF4 quiescence contrast — not the observation that fibroblasts express collagen, which would not be surprising.

The biological convergence merits explicit, but disciplined, interpretation. The six organ contexts integrated here are driven by upstream cues that are mechanistically heterogeneous — mechanical loading and orthodontic strain in the periodontal ligament, biofilm-driven inflammation and dysbiosis in periodontitis, hypoxia and TGF-β-rich tumor stroma in oral squamous-cell carcinoma and colorectal cancer, intra-articular pressure and cartilage-derived DAMPs in temporomandibular-joint osteoarthritis, and cholestatic injury in bile-duct-ligated liver. Yet all six converge on a common POSTN-amplified ECM-remodelling output. The Lei et al. mechanistic dissection in liver portal fibroblasts[3] suggests one parsimonious model for this convergence: KLF4 acts as a transcriptional brake that directly represses *POSTN* in quiescent portal fibroblasts, and POSTN — once derepressed — engages αvβ5 integrin to amplify activation in an autocrine and paracrine loop. We propose, but do not demonstrate here, that an analogous KLF4 → POSTN reciprocal mechanism is compatible with the transcriptional pattern we observe in the other five contexts. The cross-tissue transcriptional conservation (102-gene shared core, 6/6 contexts upregulating POSTN itself) is consistent with — but does not on its own establish — a convergent activation node operating across tissues. Treating this as a testable model rather than a demonstrated pathway, the predictions are: (i) tissue-specific upstream cues (mechanical strain, hypoxia, TGF-β, cholestasis) should each be sufficient to lift KLF4-mediated *POSTN* repression in the relevant fibroblast compartment; (ii) loss of KLF4 in resting non-hepatic fibroblasts should be sufficient to shift cells toward POSTN+ states; (iii) αvβ5 blockade should attenuate POSTN-driven activation in non-hepatic fibroblasts. Functional confirmation — KLF4 perturbation, αvβ5 blockade, or live-cell autocrine/paracrine assays in non-liver contexts — remains future work.

As a sequence-level companion to the mouse functional evidence, we asked whether the human *POSTN* promoter is at least sequence-compatible with KLF4 binding. We scanned the human *POSTN* promoter (−2 kb to +0.5 kb of the TSS, GRCh38) for canonical KLF4 motifs (JASPAR MA0039.4) at a relative-score cutoff of 0.75 (Fig. S2) and identified four hits — two clustered at the TSS (relative score 0.85 at +231 bp on the minus strand, AGGGGTCGGGAC; 0.82 at +32 bp on the plus strand, GTTCCCTCCCAC) and two upstream (−996 and −1,764 bp). At a relative-score cutoff of 0.75, KLF4 motif matches are common in mammalian DNA — under a uniform background a 12-bp PSSM yields on the order of 18 expected hits per 2.5-kb both-strand window — so the four observed hits in the human *POSTN* promoter constitute sequence compatibility, not enrichment evidence. We therefore interpret Fig. S2 as showing that the human *POSTN* proximal promoter contains KLF4-like motifs at conventional FIMO-style scoring stringency (consistent with prior mouse ChIP-qPCR and luciferase-reporter data[3]), but not as independent evidence of human KLF4 binding. Direct experimental validation of KLF4 occupancy on the human *POSTN* promoter remains future work.

The cluster-consensus POSTN+ definition trades sensitivity for specificity by design: a cell is POSTN+ only if its atlas neighbourhood is also POSTN-enriched and the cell itself carries a non-zero read. The resulting set is a strict subset of broader expression-based pools (recall 71–85% relative to definitions B and C; Table S1), with the largest discrepancy in periodontitis, where dispersed POSTN expression in non-candidate clusters reduces our call rate. We retain this definition as primary because it is auditable and well-anchored to the cluster structure on which the integration is built. Importantly, the primary biological conclusion of this work — recurrence of a POSTN-amplified ECM-remodelling core across all six contexts — is not load-bearing on the cell-count differences between definition A and the broader definitions B and C; the shared core gene set and the per-context POSTN+ frequency ranking survive under all three definitions (per-context Spearman ρ ≥ 0.71 for every pairwise comparison; Table S1), so the choice of A trades estimated POSTN+ pool size for auditability rather than altering which programme is conserved across tissues.

Several limitations should be acknowledged. To evaluate whether the cross-tissue POSTN+ structure depends on the integration method, we reproduced the atlas with scVI[18] (a deep generative model) on the seven datasets distributed with raw counts, using the 3,000 highly variable genes identified by a batch-aware seurat_v3 selection (Fig. S3; results/integration_benchmark.tsv). Global batch-mixing entropy was comparable between scVI (0.949, 33.8% of the seven-batch theoretical maximum log2(7) = 2.807) and Harmony at matched input (0.860, 30.6%), and was substantially above the eight-dataset primary Harmony entropy (0.303, 10.1% of log2(8) = 3.000). The eight-dataset entropy is therefore primarily explained by the inclusion of the pre-scaled perio_mmp_gse171213_caf cohort, which has no raw counts and clusters apart from the rest under either method, rather than by Harmony-specific under-correction.

Decomposing the cluster-consensus POSTN+ definition into its two criteria across the primary 8-set Harmony, the method-matched Harmony 7-set 3k-HVG, and the scVI embedding reveals a more specific finding (Fig. S3C,D; results/drift_decomp_per_ctx.tsv). Criterion (b) — per-cell POSTN expression positivity in log1p-normalized counts — is method-invariant in all six contexts: TMJ ≈ 48.6%, liver ≈ 62.1%, OSCC ≈ 54.1%, periodontitis ≈ 50.9%, CRC ≈ 42.6%, and PDL ≈ 17.7% under every integration configuration. The drift in per-context POSTN+ frequencies under method change therefore lives entirely in criterion (a), the cluster-level Wilcoxon padj < 0.05 and log2FC > 0.5 filter. In the primary 8-set Harmony embedding, mouse cells form a near-pure mega-cluster (leiden cluster 23: 99.5% mouse, 82% liver, 7,883 cells) within which POSTN is markedly elevated relative to the rest of the atlas, and the cluster passes criterion (a) — so 86% of liver cells and 72% of TMJ cells are flagged as in candidate clusters. Under better-mixed embeddings the same mouse cells are dispersed across multiple mixed-composition clusters (scVI places only 45% of mouse cells in mouse-majority clusters versus 86–88% under Harmony at any HVG configuration), the cluster-level POSTN signal in those mixed clusters falls below threshold, and criterion (a) becomes more conservative. The 8-set Harmony’s high mouse-context POSTN+ frequencies (TMJ 40.8%, liver 55.1%) therefore reflect a specific interaction between the conservative cluster-consensus definition and the under-correction of mouse data, not a stable cross-tissue biological prevalence. We characterise the per-context POSTN+ frequencies as definition-A estimates under the primary Harmony embedding rather than as integration-invariant biological percentages, and emphasise that the load-bearing claims of this study — the cross-tissue recurrence of POSTN+ cells (every context contains POSTN+ cells under every method tested), the 102-gene shared core program (recovered in human-only re-derivation; Table S3), and the POSTN+/KLF4 contrast — are not load-bearing on the absolute mouse-context POSTN+ frequencies. scANVI[19] was not run because no harmonised cell-type label is available across the eight cohorts; the only globally defined cell-level annotation in our data (tissue_context) is highly collinear with the integration batch and would therefore bias the semi-supervised step toward batch retention rather than batch correction, and we defer scANVI to a later revision when atlas-level marker-based cell-type annotation will provide an unbiased label set.

Two further limitations are unchanged from the M1 protocol. The MMP-CAF periodontitis dataset was distributed only in normalized form; sensitivity analysis 3 demonstrates that this cohort does not drive the shared core, and the cross-method decomposition above further isolates it as the only source of the conservative eight-dataset entropy. Full cross-validation against the raw-count-only atlases of Buechler 2021[1] and Korsunsky 2022[2] is future work. The Puram 2017 OSCC cohort was treated as a supplementary validation rather than included in the main atlas because TPM-only data are not compatible with the raw-count-aware Harmony pipeline. Statistical power for context-level comparisons is limited by n = 6 contexts; we report Spearman correlations together with exact permutation tests rather than asymptotic p-values for this reason.

The atlas defines POSTN+ states transcriptionally and provides a substrate for the next layer of analysis. Spatial atlases of POSTN+ stroma have begun to appear, including a recent rectal-cancer Xenium panel[20] that operates at spatial resolution rather than the integration level; our atlas and these spatial works are complementary rather than competing. Lineage tracing, niche-level organization, and therapeutic perturbation prediction on the POSTN+ cluster — including the KLF4 → POSTN axis as a candidate druggable contrast — are forecast as next-stage work. The 102-gene shared core program and its cross-tissue conservation provide a defined target list for that perturbation work, and the POSTN+ versus KLF4-quiescent contrast provides the mechanistic axis along which interventions can be evaluated. Functional validation of the KLF4-POSTN regulatory axis in human oral fibroblasts is ongoing and will be reported separately.

## Methods

### Data sources

Eight core datasets covering six organ contexts were integrated (Table 1; full per-dataset metadata, including file paths and md5 checksums, is provided in `data/manifest.tsv` of the project repository).

**Table 1.**
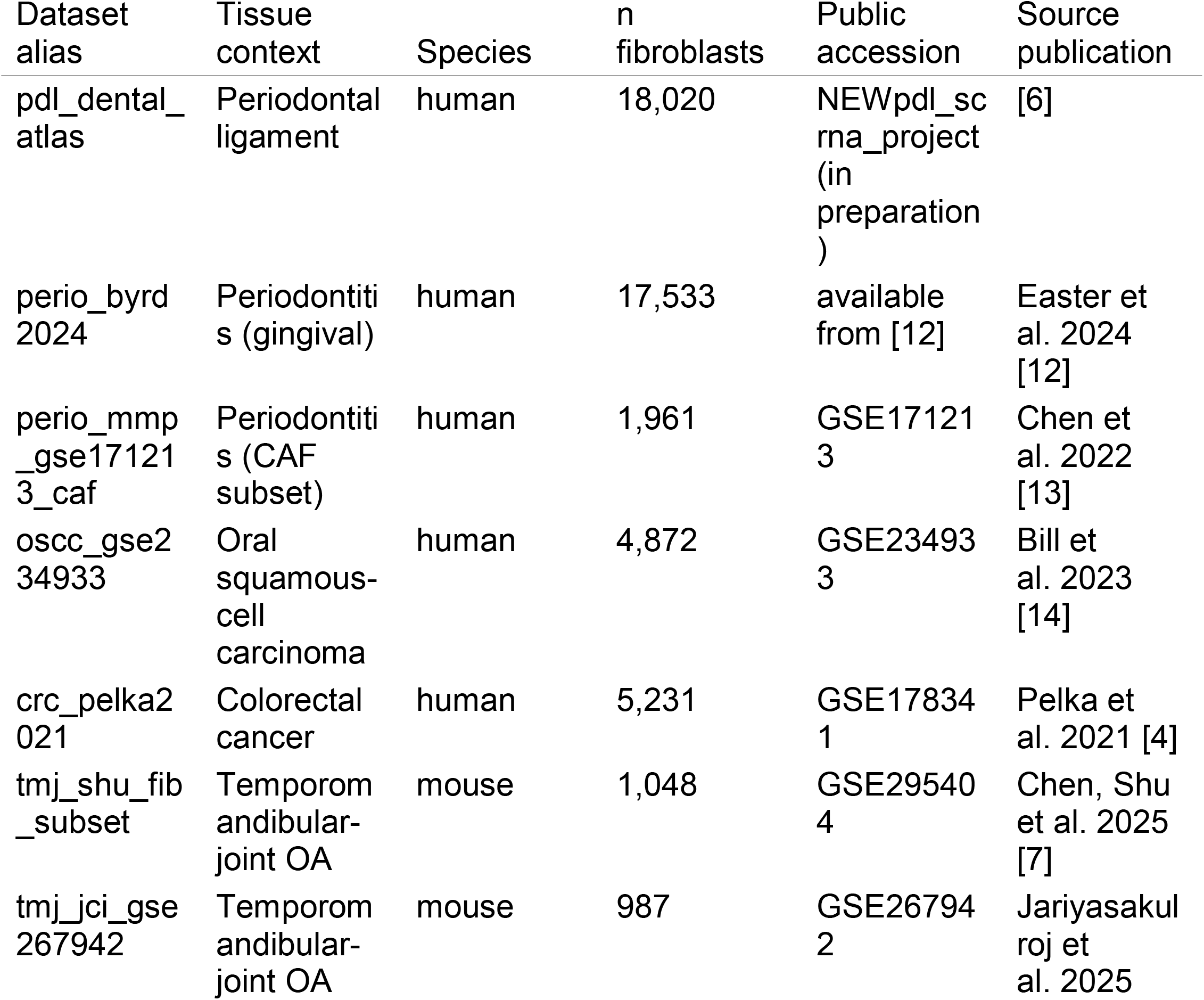
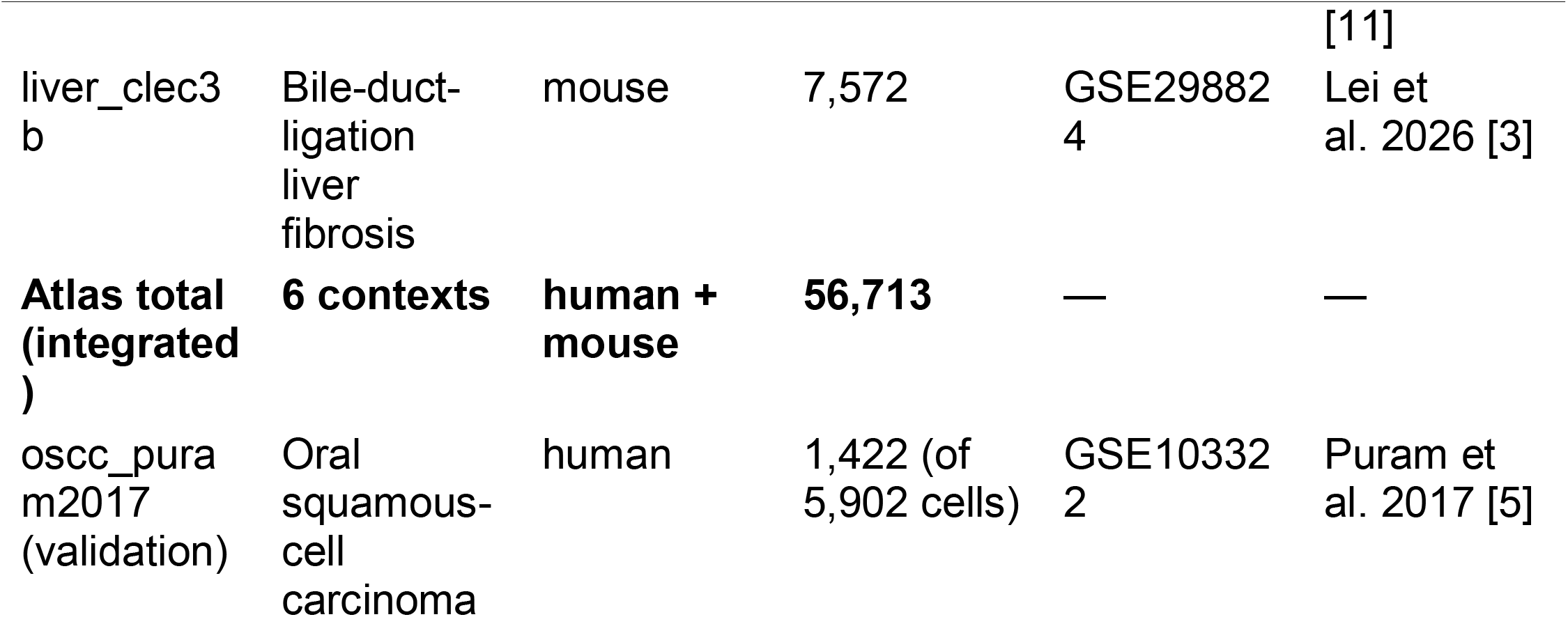
Datasets integrated into the cross-tissue POSTN+ atlas.

The Puram et al. 2017 head-and-neck cohort (GSE103322; 5,902 cells, of which 1,422 are author-annotated fibroblasts) served as an independent supplementary validation cohort outside the main integration because it is distributed as TPM rather than raw counts and is therefore incompatible with the raw-count-aware Harmony pipeline (last row of Table 1).

### Per-dataset pre-processing

Published cell-type assignments were respected for datasets where they were available, and we did not re-run author-level QC except where strictly necessary. The liver bile-duct-ligation cohort required per-sample QC (min_genes between 200 and 7,000; pct_mt < 20) and a leiden clustering at resolution 0.5 to identify fibroblast clusters by the co-presence of Col1a-family genes and Pdgfra in the top fifty markers; clusters 1 and 22 met this criterion (7,572 cells), and cluster 0 — bearing Reln, Cxcl12, and other hepatic-stellate-cell markers — was explicitly excluded to match the published distinction between Clec3b+ portal fibroblasts and stellate cells[3]. The Pelka colorectal-cancer cohort was prepared by reading the deposited GSE178341 .h5 matrix, merging the published cluster.csv and metatables.csv into the .obs slot, and subsetting to clMidwayPr == ‘Fibro’ (5,231 cells; eleven CAF subtypes preserved). The MMP-CAF periodontitis cohort was distributed only as a normalized expression matrix; we retained it for Harmony integration but excluded it from analyses that require raw counts (most importantly, the per-context shared-core derivation; see also the Sensitivity Analyses section).

### Cross-species ortholog mapping

The mouse cohorts (TMJ shu, TMJ jci, and liver_clec3b) were re-indexed from mouse to human gene symbols using a 1:1 ortholog table built from MGI HOM_MouseHumanSequence.rpt (downloaded 2026-05-07). We retained 18,782 1:1 pairs from 21,930 mouse and 24,592 human entries; one-to-many and many-to-many relationships were discarded to avoid downstream ambiguity. POSTN-axis markers (POSTN, COL1A1/A2, SPARC, KLF4, CLEC3B, ACTA2) all mapped 1:1, and unmapped genes — predominantly Gm-prefix predicted genes without human orthologs — were dropped before concatenation. The full ortholog table is provided as `results/ortholog_map.tsv.`

### Atlas integration

The eight datasets were concatenated on the intersection of expressed genes (14,545 genes; min-cells = 10 filter retained 14,296 genes), normalized per cell to a target sum of 10^4^ counts, and log1p-transformed. PCA was performed on the full filtered gene set rather than on a high-variable-gene subset; we did not subset to HVGs because both the seurat and seurat_v3 flavours of scanpy.pp.highly_variable_genes produced numerically degenerate results on this matrix (a pandas-NA propagation issue with seurat and a LOESS extrapolation on extreme means with seurat_v3). Fifty PCA components were used as input to Harmony[15], which was called via harmonypy 0.2.0 directly rather than through scanpy.external.pp.harmony_integrate (the wrapper exhibited a shape inconsistency on our input matrix). Harmony parameters were the package defaults (θ = 2, σ = 0.1, max_iter = 20, batch_key = dataset_alias) and the algorithm converged in five iterations. A k = 15 nearest-neighbour graph was built on the Harmony embedding, and a leiden clustering at resolution 0.5 (igraph backend, two iterations, undirected graph) yielded 56 clusters which serve as the cluster substrate for the POSTN+ definition.

### Cluster-level differential expression

Wilcoxon rank-sum differential expression was computed via scanpy 1.12 rank_genes_groups (use_raw = False, on log1p-normalized X). The logfoldchanges field returned by scanpy was numerically NaN on this matrix (a known issue in some sparse/categorical configurations), and we replaced it with manually computed log2((mean_in + 1) / (mean_out + 1)) on raw counts where mean_in is the per-gene mean count among cells in the focal cluster and mean_out the mean among all other cells. The pseudocount of 1 was chosen for robustness against dropout-driven noise.

### POSTN+ cell definition

The POSTN+ definition is cell-level and combines two conditions, both of which must hold:

a. the cell’s atlas-integrated leiden cluster has POSTN with Wilcoxon padj < 0.05 and manually computed log2FC > 0.5 against all other clusters; and
b. the cell’s normalized POSTN expression is greater than zero.

Five clusters met condition (a): 14, 16, 17, 22, and 23. Cells from these clusters with non-zero normalized POSTN expression numbered 11,451 (20.2% of the atlas). The cluster condition acts as a consensus filter that excludes high-POSTN expressors from non-enriched clusters; alternatives are evaluated in the Sensitivity Analyses section.

### Shared core program derivation

Within each context, mean raw counts were computed in POSTN+ and POSTN-fibroblasts, and per-gene log2FC was computed as log2((mean_pos + 1) / (mean_neg + 1)). A gene was retained for the per-context up-regulated set if its log2FC exceeded 1.0 and the mean POSTN+ count exceeded 0.5 raw counts (the latter filtering against dropout-driven false positives). Across the six contexts, a gene was retained in the shared core program if it was upregulated by these criteria in at least four of the six contexts. This produced 102 genes. The full pergene per-context log2FC table, including all genes tested and their up-context counts, is provided as `results/shared_core_program.tsv` .

### Sensitivity analyses

Three sensitivity analyses were pre-registered and executed before manuscript drafting.

#### Harmony parameter (θ)

The atlas integration was re-run with θ = 4, σ = 0.05, max_iter = 30. The mean cluster-level dataset entropy rose from 0.303 to 0.593 (max 3.000 for eight batches), the leiden clustering at resolution 0.5 yielded 37 clusters (vs 56 at default θ), six clusters met the cluster-level POSTN criterion (vs five), and 10,608 cells (vs 11,451) were classified POSTN+. Per-context POSTN+ frequencies shifted by less than 4.4 percentage points absolute, and the per-context Spearman rank correlation with the default-θ frequencies was ρ = 1.000 (n = 6 contexts; exact permutation p = 0.0028). The overlap of the top twenty-five shared-core genes between parameterizations was 18/25 (72%). The full per-context table and top-25 gene overlap are provided as Table S2 (raw data: `results/sensitivity_harmony_theta4.tsv`).

#### POSTN+ definition (A/B/C)

Three definitions were evaluated on the same atlas:(A) the cluster-consensus definition above (n = 11,451; 20.2%); (B) a percentile threshold of POSTN raw counts ≥ 70th percentile, requiring at least one read (n = 17,948; 31.6%); and (C) a six-gene module score (POSTN + COL1A1 + SPARC + BGN + FN1 + TGFB1) thresholded at the top 30% (n = 17,014; 30.0%). Recall of A within B was 85.4%, A within C 71.2%, and B within C 73.7%. Per-context Spearman rank correlations were ρ = 0.714 (A vs B), 0.886 (A vs C), and 0.943 (B vs C); n = 6 contexts; exact permutation. The full pairwise overlap and per-context frequency table is provided as Table S1 (raw data: `results/sensitivity_postn_definition.tsv`).

#### Dataset exclusion (no MMP-CAF)

The integration was re-run after excluding perio_mmp_gse171213_caf (the only normalized-only cohort, n = 1,961). The re-derivation produced 105 shared core genes, of which 100 of the original 102 were retained (98% overlap). All twenty-five a-priori expected ECM markers were preserved. The full gene-level table is provided as `results/shared_core_no_mmp_caf.tsv` .

#### Cross-species robustness (human-only re-derivation)

The integrated atlas was restricted to the five human datasets (47,106 cells across four contexts: PDL, perio, OSCC, CRC) and the per-context POSTN+/POSTN-shared core was re-derived under the same per-gene criterion (log2FC > 1.0 AND mean POSTN+ raw counts ≥ 0.5) at a ≥ 3-of-4 (75%) cross-context threshold (parallel to the baseline 4-of-6 = 67%). All 102 baseline shared core genes were recovered (Table S3; raw data: `results/shared_core_human_only.tsv`); the human-only set contained 238 genes in total (the 102 baseline genes plus 136 additional genes admitted by the more-permissive 75% threshold). POSTN itself remained upregulated in 4/4 human contexts.

### In-silico KLF4 binding-motif scan of the human POSTN promoter

The human *POSTN* gene (Ensembl ENSG00000133110; chromosome 13, − strand) was queried via the Ensembl REST `/lookup/symbol`endpoint, and the −2,000 bp to +500 bp promoter region around the TSS (GRCh38, chr13:37,598,355–37,600,855) was retrieved as the sense strand via `/sequence/region` . The KLF4 position frequency matrix (JASPAR MA0039.4, length 12 bp) was retrieved from the JASPAR REST API and converted to a position-specific scoring matrix with pseudocount 0.5 under a uniform 0.25 background. The promoter was scanned on both strands using `Bio.motifs.search`(Biopython 1.84) at a relative-score cutoff of 0.75. Hits were tabulated with their position relative to the TSS, strand, matched k-mer (reported as the sense-strand sequence), absolute score, and relative score, and rendered as Figure S2 (raw data: `results/klf4_motif_hits_postn_promoter.tsv`). We note explicitly that a relative-score cutoff of 0.75 on a 12-bp PSSM is a conventional FIMO-style cut and is permissive at the genome scale: under a uniform 0.25 background, the same cutoff yields on the order of 0.0037 expected hits per 12-mer per strand, ≈ 18 expected hits per 2.5-kb both-strand window, and on the order of 23 million expected hits across the 3.1-Gb human genome. The motif scan is therefore reported as a sequence-compatibility check — the human *POSTN* promoter contains KLF4-like motifs at conventional scoring stringency — not as an enrichment or in-vivo binding test.

### Independent validation (Puram 2017 cohort)

The GSE103322 TPM matrix was loaded with metadata rows separated from gene-expression rows by a numeric-row test. Gene names in this cohort are wrapped in literal single quotes and were stripped before symbol matching. Of 5,902 cells, 1,422 were annotated by Puram and colleagues as fibroblasts under the cohort’s own non-cancer cell-type column; 1,112 were primary-tumour and 310 were lymph-node-metastatic fibroblasts. The atlas-derived 102-gene shared core program was scored per cell with scanpy.tl.score_genes (random_state = 0). 101 of 102 core genes were present in the Puram TPM matrix. The mean core score in lymph-node-metastatic fibroblasts (2.85) exceeded that in primary fibroblasts (2.76) at Mann–Whitney U p = 0.0046 (Welch’s t-test p = 0.049). The full sidecar is provided as `figures/FigS1_puram_validation.json` .

### Statistical reporting and reproducibility

Spearman rank correlations across n = 6 contexts are reported with exact permutation p-values rather than asymptotic approximations, in keeping with reviewer recommendation that small-n rank statistics not be reported as p = 0. Each manuscript figure ships with a JSON sidecar listing every cited number, color palette, and reproduce command (CLAUDE.md Section 3.5). All cited numbers in this manuscript are registered in `number_provenance.md` . Analysis code will be released in a public repository upon journal acceptance. During peer review, code is available from the corresponding author upon reasonable request. Scripts are organized as follows: 00_* (data ingestion and ortholog mapping), 02_* and 03_* (concatenation and Harmony integration), 04_* (POSTN definition application), 05_* (main figure), 06a–c_* (sensitivity analyses), and 07_* (Puram validation). The analysis environment is the bindeath conda environment (scanpy 1.12, anndata, harmonypy 0.2.0, R 4.3.3, Seurat 5.4). Processed atlas .h5ad files and the ortholog map will be deposited at Zenodo upon journal acceptance; raw datasets are available under the GEO and other accessions listed in Table 1.

## Data and Code Availability

All raw datasets are available under the public accessions listed in Table 1 (GEO GSE171213, GSE234933, GSE267942, GSE295404, GSE298824, GSE178341, and GSE103322; the periodontal-ligament cohort and the Byrd-2024 periodontitis cohort are available from their respective deposits cited in [6, 12]). The processed integrated atlas (.h5ad), the 102-gene shared core program, the three sensitivity-analysis tables (Tables S1, S2, and the no-MMP-CAF re-derivation), and the per-figure JSON sidecar registry that support the findings of this study are available from the corresponding author (Guoli Yang) upon reasonable request, and will additionally be deposited at Zenodo upon journal acceptance; the DOI will be included in the final published version.

## Author Contributions

C.W. designed the study, performed all data integration and analyses, and drafted the manuscript. H.Y., M.L., and Y.W. contributed to data curation and manuscript revision. G.Y. supervised the project, secured funding, and revised the manuscript. All authors read and approved the final version.

## Acknowledgments

We thank the contributing investigators of the GSE103322, GSE171213, GSE178341, GSE234933, GSE267942, GSE295404, and GSE298824 cohorts for making their primary data publicly available, without which a cross-tissue integration of this scope would not be possible.

## Ethics Statement

This study analyzes only publicly deposited single-cell datasets; no new human or animal experiments were performed.

---

**Figure S1.**
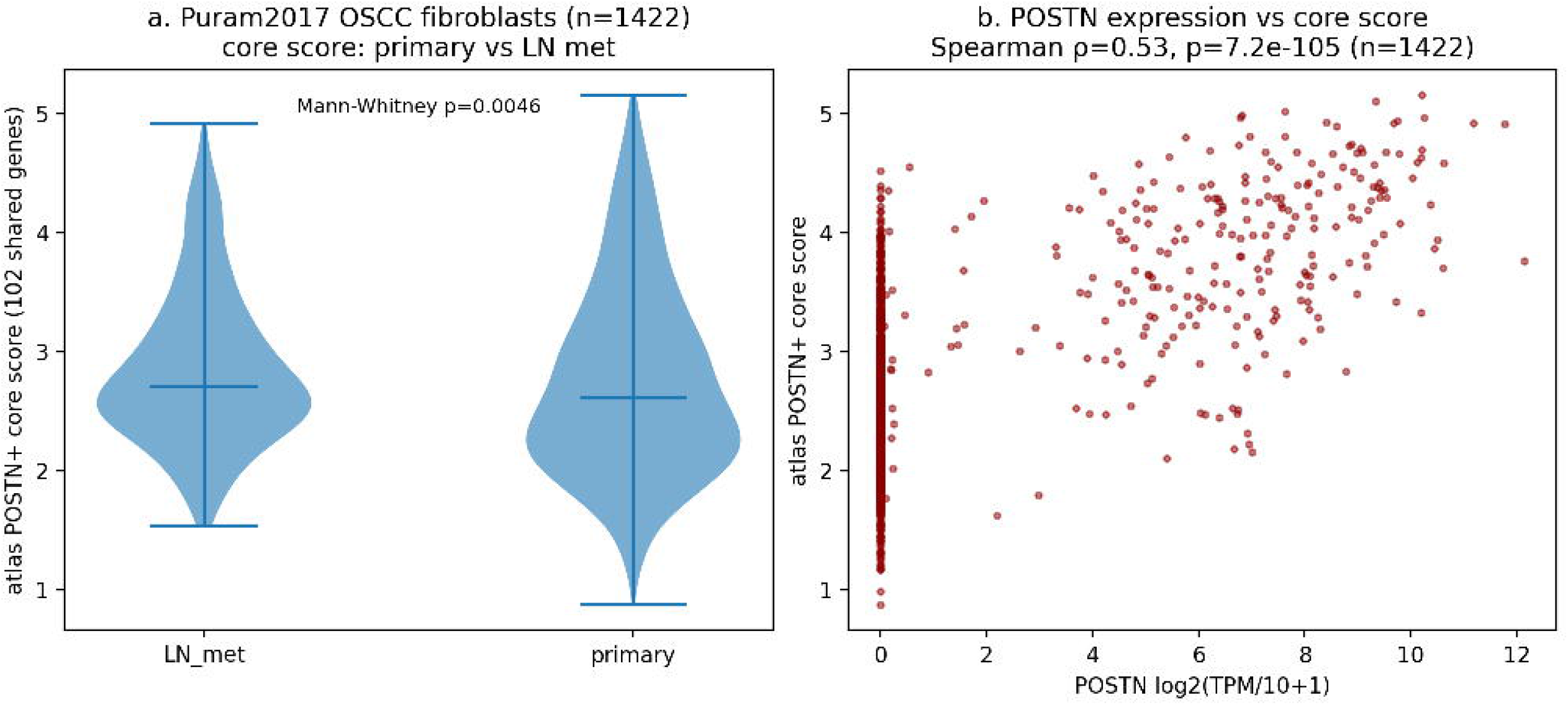
Independent validation in the Puram 2017 OSCC cohort. Left: violin distributions of the atlas-derived 102-gene core score per cell among Puram-annotated fibroblasts, split by primary tumour (n = 1,112) versus lymph-node metastasis (n = 310); Mann–Whitney U p = 0.0046. Right: per-cell POSTN log2(TPM/10 + 1) versus core score (Spearman correlation reported in the title).

**Figure S2.**
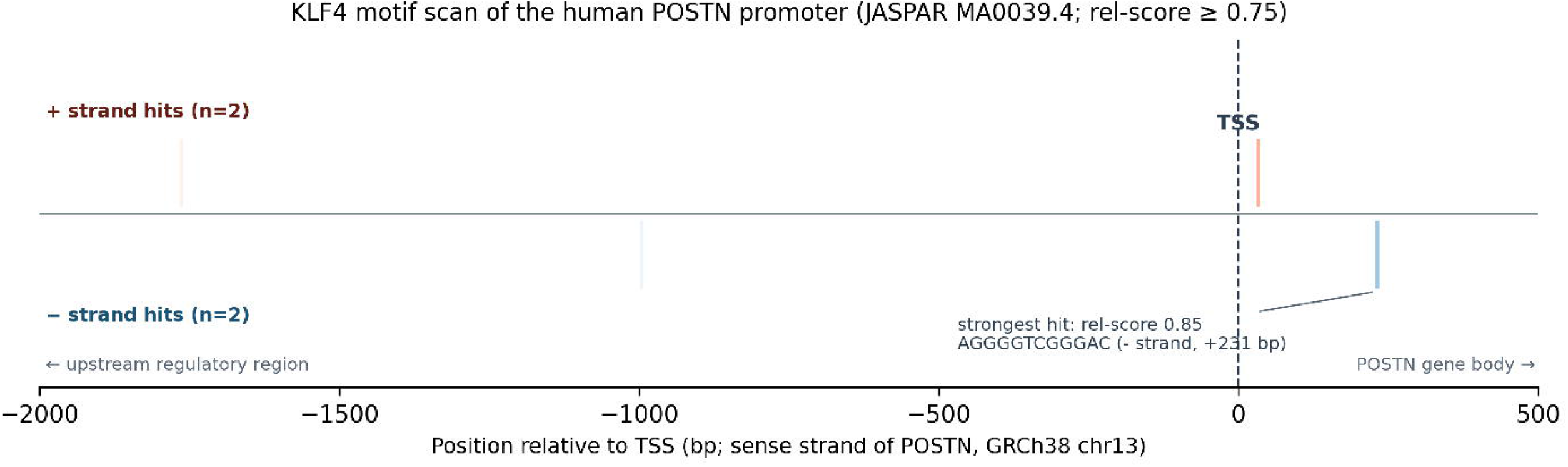
In-silico KLF4 binding-motif scan of the human POSTN promoter (JASPAR MA0039.4, GRCh38). Single-track motif ideogram of the −2 kb to +0.5 kb promoter region around the *POSTN* transcription start site, with KLF4 motif hits (relative score ≥ 0.75) drawn as vertical ticks above (+ strand, n = 2) and below (− strand, n = 2) the promoter axis; tick colour scales with relative score. The strongest hit (rel-score 0.85, AGGGGTCGGGAC, − strand, +231 bp) is annotated. The figure is shown as a sequence-compatibility check: at relative score ≥ 0.75 the 12-bp KLF4 PSSM is expected to yield on the order of 18 hits per 2.5-kb both-strand window under a uniform background, so the four observed hits are consistent with KLF4 motifs being *present* at the human *POSTN* proximal promoter at conventional FIMO-style stringency, but not with statistical enrichment. The figure complements the direct ChIP-qPCR and luciferase-reporter evidence reported in mouse portal fibroblasts[3]; experimental confirmation of KLF4 occupancy on the human *POSTN* promoter remains future work.

**Figure S3.**
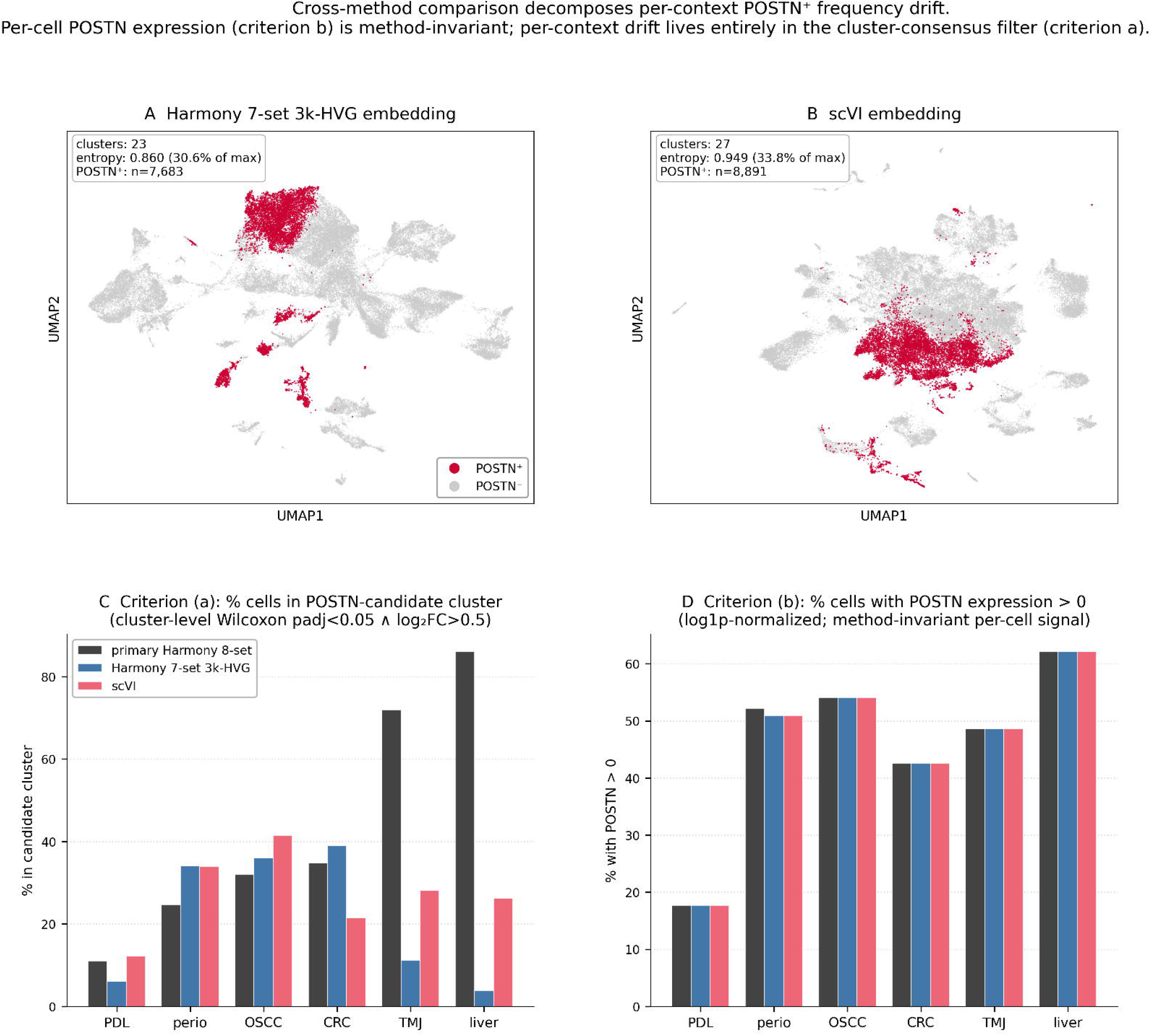
Cross-method comparison decomposes per-context POSTN+ frequency drift into a method-invariant per-cell expression signal and a method-dependent cluster-consensus filter. **A**. UMAP of the Harmony 7-set 3k-HVG embedding (55,263 fibroblasts; mean cluster-level dataset entropy 0.860, 30.6% of log2(7) = 2.807; 23 leiden clusters at resolution 0.5; n = 7,683 POSTN+ cells highlighted in red on grey). **B**. UMAP of the scVI embedding on the same seven datasets and the same 3,000 batch-aware highly variable genes (entropy 0.949, 33.8% of maximum; 27 clusters; n = 8,891 POSTN+ cells in red). The perio_mmp_gse171213_caf cohort is excluded from both panels because it is distributed only as scaled values (minimum −3.7), incompatible with scVI’s negative-binomial likelihood; periodontitis context is still covered by perio_byrd2024 (17,533 cells). **C**. Per-context grouped bars for criterion (a) of the POSTN+ definition — the fraction of cells whose atlas leiden cluster passes Wilcoxon padj < 0.05 and log2FC > 0.5 for POSTN — across the primary 8-set Harmony atlas, the method-matched Harmony 7-set 3k-HVG embedding, and scVI. The cluster-level criterion drifts dramatically in the mouse contexts (liver 86.0% → 3.8% → 26.2%; TMJ 71.9% → 11.2% → 28.1%) but is stable in human contexts (PDL 11.0 → 6.1 → 12.2%; perio 24.7 → 34.0 → 34.0%; OSCC 32.0 → 36.0 → 41.5%; CRC 34.7 → 39.0 → 21.5%). The mouse-context drift tracks mouse-cell cluster purity: 86.4% of mouse cells live in mouse-majority clusters under primary 8-set Harmony (driven by the near-pure mouse mega-cluster leiden #23, 99.5% mouse / 82% liver / 7,883 cells), 88.0% under Harmony 7-set 3k-HVG, but only 45.2% under scVI. **D**. The corresponding per-context grouped bars for criterion (b) — the fraction of cells with POSTN expression > 0 in log1p-normalized counts — are visually flat across all three integration configurations (PDL 17.7%; perio 51–52%; OSCC 54.1%; CRC 42.6%; TMJ 48.6%; liver 62.1%). The decomposition therefore demonstrates that per-cell POSTN expression positivity is method-invariant, and the per-context drift observed in the final POSTN+ frequencies (Fig. 1B right; Discussion) is entirely attributable to the cluster-consensus filter.

