## Supplementary material for "A cross-tissue POSTN+ fibroblast atlas links periodontal, tumor, and fibrotic stromal niches": Table_S1-S3: Supplementary_Information.docx

### Supplementary Tables

**Table S1. POSTN⁺ definition robustness (A vs B vs C).** Three POSTN⁺ definitions evaluated on the integrated 56,713-cell atlas. **A** = §0 cluster-consensus (cell in candidate cluster meeting Wilcoxon padj < 0.05 ∧ log₂FC > 0.5 AND POSTN expression > 0; primary definition). **B** = raw POSTN counts at the 70th percentile (≥ 1 read). **C** = six-gene module score (POSTN + COL1A1 + SPARC + BGN + FN1 + TGFB1) in the top 30%.

*S1a — Definition sizes.*

| Metric | A (cluster-consensus) | B (percentile 70) | C (module score) |
| --- | --- | --- | --- |
| n cells classified POSTN⁺ | 11,451 | 17,948 | 17,014 |
| % of atlas (n = 56,713) | 20.2 | 31.6 | 30.0 |

*S1b — Per-context POSTN⁺ frequency (%, sorted by definition A).*

| Context | A | B | C |
| --- | --- | --- | --- |
| PDL | 6.2 | 9.8 | 5.8 |
| perio | 16.4 | 40.6 | 34.9 |
| CRC | 19.5 | 34.9 | 13.5 |
| OSCC | 24.7 | 45.8 | 60.3 |
| TMJ | 40.8 | 39.6 | 53.2 |
| liver | 55.1 | 47.5 | 60.9 |

*S1c — Pairwise overlap and per-context rank concordance (n = 6 contexts; exact permutation).*

| Pair | ∩ cells | Jaccard | %first ∈ second | %second ∈ first | Spearman ρ | exact p |
| --- | --- | --- | --- | --- | --- | --- |
| A vs B | 9,779 | 0.498 | 85.4 | 54.5 | 0.714 | 0.136 |
| A vs C | 8,148 | 0.401 | 71.2 | 47.9 | 0.886 | 0.033 |
| B vs C | 12,541 | 0.559 | 69.9 | 73.7 | 0.943 | 0.017 |

*S1d — Triple agreement.* Cells classified POSTN⁺ by all three definitions: **7,486 (13.2% of atlas)**; cells classified POSTN⁺ by at least one definition: 23,431 (41.3%); within-union consensus rate 31.9%.

**Table S2. Harmony θ-parameter sensitivity (default θ = 2 vs aggressive θ = 4).** Same eight datasets re-integrated under more aggressive batch correction (θ = 4, σ = 0.05, max_iter = 30 vs default θ = 2, σ = 0.1, max_iter = 20).

*S2a — Global integration metrics.*

| Metric | θ = 2 (default) | θ = 4 (sensitivity) |
| --- | --- | --- |
| Leiden clusters at res 0.5 | 56 | 37 |
| POSTN⁺ cells (n; %) | 11,451 (20.2%) | 10,608 (18.7%) |
| Candidate clusters (a-condition) | 5 (14, 16, 17, 22, 23) | 6 (11, 14, 15, 19, 21, 23) |
| Mean cluster dataset entropy | 0.303 | 0.593 |
| Spearman ρ on per-context POSTN⁺ % rank (n = 6) | — | 1.000 (exact permutation p = 0.0028) |
| Top-25 shared-core gene overlap | — | 18 / 25 (72%) |

*S2b — Per-context POSTN⁺ frequency (sorted by θ = 2 frequency).*

| Context | n cells | θ = 2 (%) | θ = 4 (%) | Δ (pp) |
| --- | --- | --- | --- | --- |
| PDL | 18,020 | 6.2 | 5.3 | −0.9 |
| perio | 18,983 | 16.4 | 15.1 | −1.3 |
| CRC | 5,231 | 19.5 | 18.3 | −1.2 |
| OSCC | 4,872 | 24.7 | 23.1 | −1.6 |
| TMJ | 2,035 | 40.8 | 36.4 | −4.4 |
| liver | 7,572 | 55.1 | 52.4 | −2.7 |

*S2c — Top-25 shared-core gene overlap.* Common to both θ values (n = 18): AEBP1, BGN, CALD1, COL11A1, COL12A1, COL1A1, COL1A2, COL3A1, COL5A1, COL5A2, COL6A1, COL6A2, COL6A3, CTHRC1, CTSK, LGALS1, MMP2, POSTN. Only at θ = 2: CD63, CDH11, PALLD, PCOLCE, PPIB, RPL15, RPS2. Only at θ = 4: FN1, IGFBP5, PMEPA1, SERPINH1, SPARC, TPM4, VCAN. The non-overlap is dominated by re-ordering of additional ECM markers (FN1, SPARC, SERPINH1) into and out of the top-25 cut, not by gain or loss of distinct biology.

**Table S3. Cross-species robustness — human-only re-derivation of the shared core program.** The integrated atlas was restricted to the five human datasets (47,106 cells across PDL, perio, OSCC, CRC) and the per-context shared core was re-derived under the same per-gene retention criterion at a parallel ≥ 3-of-4 (75%) threshold.

*S3a — Atlas restriction.*

| Metric | Baseline (6 contexts, human + mouse) | Human-only (4 contexts) |
| --- | --- | --- |
| Cells | 56,713 | 47,106 |
| Datasets | 8 | 5 |
| Threshold for shared-core retention | ≥ 4/6 (67%) | ≥ 3/4 (75%) |
| Shared-core genes | 102 | 238 |
| Baseline genes recovered in human-only set | — | 102 / 102 (100%) |
| Genes lost when dropping mouse cohorts | — | 0 |

*S3b — POSTN per-context log₂FC in the human-only re-derivation (POSTN⁺ vs POSTN⁻).*

| Context | log₂FC |
| --- | --- |
| PDL | 4.75 |
| OSCC | 2.80 |
| CRC | 3.14 |
| perio | 2.40 |

*S3c — Status of the 25 a-priori expected ECM markers in the human-only shared core.* All twenty-five markers are present, with twenty-three retained at 4/4 contexts upregulated (POSTN, COL1A1, COL1A2, COL3A1, COL5A1, COL5A2, COL6A1, COL6A2, COL6A3, COL11A1, COL12A1, SPARC, BGN, FN1, MMP2, AEBP1, CTHRC1, CALD1, LGALS1, VCAN, SERPINH1, TPM4) and two retained at 3/4 contexts (TIMP1, ASPN); FBN1 was absent from the baseline shared core and remained absent (2/4 contexts).

The baseline 102-gene shared core program is therefore recoverable from the four human contexts alone, and is not dependent on the inclusion of the mouse cohorts; the analysis cannot strictly rule out species-specific effects because dropping mouse also drops the TMJ and liver contexts entirely. full per-context, per-gene log₂FC table including all genes tested.
